# Multivariate Computational Analysis of Gamma Delta T cell Inhibitory Receptor Signatures Reveals the Divergence of Healthy and ART-Suppressed HIV+ Aging

**DOI:** 10.1101/412312

**Authors:** Anna C. Belkina, Alina Starchenko, Katherine A. Drake, Elizabeth A. Proctor, Riley M.F. Pihl, Alex Olson, Douglas A. Lauffenburger, Nina Lin, Jennifer E. Snyder-Cappione

## Abstract

Even with effective viral control, HIV-infected individuals are at a higher risk for morbidities associated with older age than the general population, and these serious non-AIDS events (SNAEs) track with plasma inflammatory and coagulation markers. The cell subsets driving inflammation in aviremic HIV infection are not yet elucidated. Also, whether ART-suppressed HIV infection causes premature induction of the inflammatory events found in uninfected elderly or if a novel inflammatory network ensues when HIV and older age co-exist is unclear. In this study we measured combinational expression of five inhibitory receptors (IRs) on seven immune cell subsets and 16 plasma markers from peripheral blood mononuclear cells (PBMC) and plasma samples, respectively, from a HIV and Aging cohort comprised of ART-suppressed HIV-infected and uninfected controls stratified by age (≤35 or ≥50 years old). For data analysis, multiple multivariate computational algorithms (cluster identification, characterization, and regression (CITRUS), partial least squares regression (PLSR), and partial least squares-discriminant analysis (PLS-DA)) were used to determine if immune parameter disparities can distinguish the subject groups and to investigate if there is a cross-impact of aviremic HIV and age on immune signatures. IR expression on gamma delta (γδ) T cells exclusively separated HIV+ subjects from controls in CITRUS analyses and secretion of inflammatory cytokines and cytotoxic mediators from γδ T cells tracked with TIGIT expression among HIV+ subjects. Also, plasma markers predicted the percentages of TIGIT+ γδ T cells in subjects with and without HIV in PSLR models, and a PLS-DA model of γδ T cell IR signatures and plasma markers significantly stratified all four of the subject groups (uninfected younger, uninfected older, HIV+ younger, and HIV+ older). These data implicate γδ T cells as an inflammatory driver in ART-suppressed HIV infection and provide evidence of distinct ‘inflamm-aging’ processes with and without ART-suppressed HIV infection.

## Introduction

Although anti-retroviral therapy (ART) has dramatically reduced the rates of HIV-associated morbidity and mortality, HIV+ individuals have a shorter lifespan than uninfected counterparts [1-3] due in part to the onset and progression of co-morbidities associated with older age, including cardiovascular disease, stroke, neurocognitive impairment, diabetes mellitus, impaired renal function, non-AIDS malignancies, and osteoporosis [4-6]. These conditions, sometimes referred to as serious non-AIDS events (SNAEs), afflict older HIV+ individuals more frequently than both younger counterparts [4, 6] and the uninfected elderly [6] and SNAE disease onset is reported to occur at younger ages in aviremic HIV+ individuals as compared with uninfected controls [5, 7], with the latter observations supporting the concept of HIV+ persons undergoing early or accelerated aging [8]. Mortality and age-associated co-morbidities among HIV+ individuals track with plasma inflammatory and coagulation markers, such as C-reactive protein (CRP), IL-6, D-dimer, and fibrinogen [9-14], similar to the general population [15-21]. These findings strongly implicate the general inflammation associated with aging, sometimes referred to as ‘inflamm-aging’ [22, 23], as an integral link to SNAE occurrence in aviremic HIV infection.

Gamma delta (γδ) T cells are a unique T cell lineage that is typically less than 10% of T cells in the circulation yet are present in considerably higher proportions in the intestinal epithelium [24-27]. With a limited T cell receptor repertoire and evidence indicating that a predominant mode of cell activation is in response to non-peptidic ligands, cytokines, and signaling via NKG2D and similar receptors [28-35], γδ T cells are implicated as early responders in immune responses and/or as innate-adaptive bridging cells, responding to innate signals to then direct the development of the adaptive immune response via secretion of cytokines and other factors. As a population, γδ T cells can exert inflammatory/cytotoxic [36, 37] or immuno-regulatory effector functions [38, 39], are integral to the control of infections [40, 41], and can exhibit potent anti-tumor activity [42-44]. The role of γδ T cells in HIV viral pathogenesis is unclear to date; shifts in γδ T cell subsets defined by T cell receptor usage (Vδ1 and Vδ2) occur early in infection [45] due to a substantial loss of the Vγ2-Jγ1.2/Vδ2+ subset that is not always recovered with viral suppression [46]. However, functional differences between γδ T cell subsets that differ in γ and δ chain usage are not well established and therefore the significance of this subset shift in relation to disease pathogenesis is uncertain. Also, circulating γδ T cells serve as a reservoir for latent HIV infection [47] and thus may have relevance in cure strategies. To date, the role of γδ T cells in the inflammation observed in aviremic HIV infection is unknown.

To understand the cellular network that drives the onset and progression of age-associated morbidities in both ART-suppressed (aviremic) HIV and healthy aging, we conducted a study that measured a large number of immune parameters from a well-characterized HIV and Aging cohort, comprised of both ART-suppressed HIV+ and age-matched uninfected controls with similar exposures and demographics, stratified by age in younger and older groups. Our results from multivariate computational analyses of high parameter single cell cytometry, plasma markers, and *ex vivo* culture supernatant cytokine data identify γδ T cells as a putative key player in the immune cell network driving ‘inflamm-aging’ in aviremic HIV infection. Also, our bioinformatic analyses revealed an novel combined impact of both virally suppressed HIV and aging on immune networks, thereby indicating that aviremic HIV+ persons do not simply prematurely age but undergo a novel inflammatory course when these two conditions collide.

## Results

Inhibitory receptor (IR) expression on γδ T cells distinguishes ART-suppressed HIV+ subjects from uninfected controls. Expression of IRs has been linked to altered functionality of immune cells [48-51]. While increased IR expression on T cell populations has been reported with aging in mice and humans [52-56], and separately with HIV infection [49, 57-59], a more comprehensive investigation of IR signatures on circulating immune cells from matched younger and older subjects with and without ART-suppressed HIV infection had not been performed to our knowledge. Therefore, in this study we analyzed PBMC from our HIV and Aging Cohort, comprised of ART-suppressed HIV+ younger (≤35yo), and older (≥50yo) subjects age-matched with uninfected counterparts (Table 1). We measured five inhibitory receptors (PD-1, TIGIT, TIM-3, CD160, LAG-3) on seven immune cell subsets (CD4+ T, CD8+ T, T regulatory (Treg), CD56^bright^ and CD56^dim^ natural killer (NK), gamma delta T (γδ T), and invariant natural killer T (iNKT) cells) using the 16-color flow cytometry panel we developed and previously described [60]. Using the CITRUS algorithm [61] we determined whether IR expression on any of the immune subsets (Supplemental Fig. 1) could be used to distinguish ART-suppressed HIV+ subjects from uninfected controls. Using 10-fold cross-validation (CV) to select the model with the minimum number of features necessary to predict these two groups, only TIGIT expression in four cellular clusters comprised of γδ T cells (Fig. 1A, clusters 1-4 in red circles), was necessary to differentiate the two subject groups with 88.6% CV accuracy (Supplemental Fig. 1). In all four clusters, TIGIT expression was higher in the ART-suppressed HIV+ subjects compared to the uninfected controls (Fig. 1B). Expression of other surface antigens on the cells in clusters 1-4 was similar for CD4 and CD127 (all negative), CD56 (all clusters intermediate) and varied for other antigens, such as CD8 (low in cluster 1, intermediate in clusters 2-4), CD16 (intermediate in clusters 1, 2, and 4, and low in cluster 3), and CD3 (all four clusters positive, with cluster 3 intermediate) (Fig. 1C). Next, a false discovery rate (FDR) threshold of 1% was used to identify all clusters that were significantly different between the two subject groups using IR expression differences. Using this method, seven clusters were significant and they all contained γδ T cells (Fig. 1D); all seven clusters had significant TIGIT expression differences between the HIV+ subjects and uninfected controls, one (cluster 3) had CD160 differences, and one (cluster 4) had TIM-3 differences. Six of the seven TIGIT expression clusters contain only or predominantly γδ T cells (Supplemental Fig. 1, Fig. 1F), with one cluster (cluster 5) containing both NK and γδ T cells; however, it is likely that γδ T cells are the sole driver of this finding given the γδ T predominance in all other clusters. Similar to the 10-fold CV results (Fig. 1B), the expression of the defining IR for clusters 3-7 was consistently higher in the HIV+ subject group compared to uninfected controls (Fig. 1E). Further phenotypic analysis of the three additional clusters that emerged from the FDR-constrained analysis (clusters 5, 6, and 7) show the presence of NK cells in cluster 5 (CD3-, CD16/56+), while clusters 6 and 7 are predominantly CD16-, CD56-, CD4-, CD8- and γδ TcR^lo^ (Fig. 1F). Together, these data demonstrate that the IR expression on circulating γδ T cells distinguishes ART-suppressed HIV+ subjects from uninfected controls. Also, the γδ T cell subsets that predict these two subject groups are diverse, including populations that differ in CD8, CD16, CD56, and CD127 expression.

**Figure 1.**
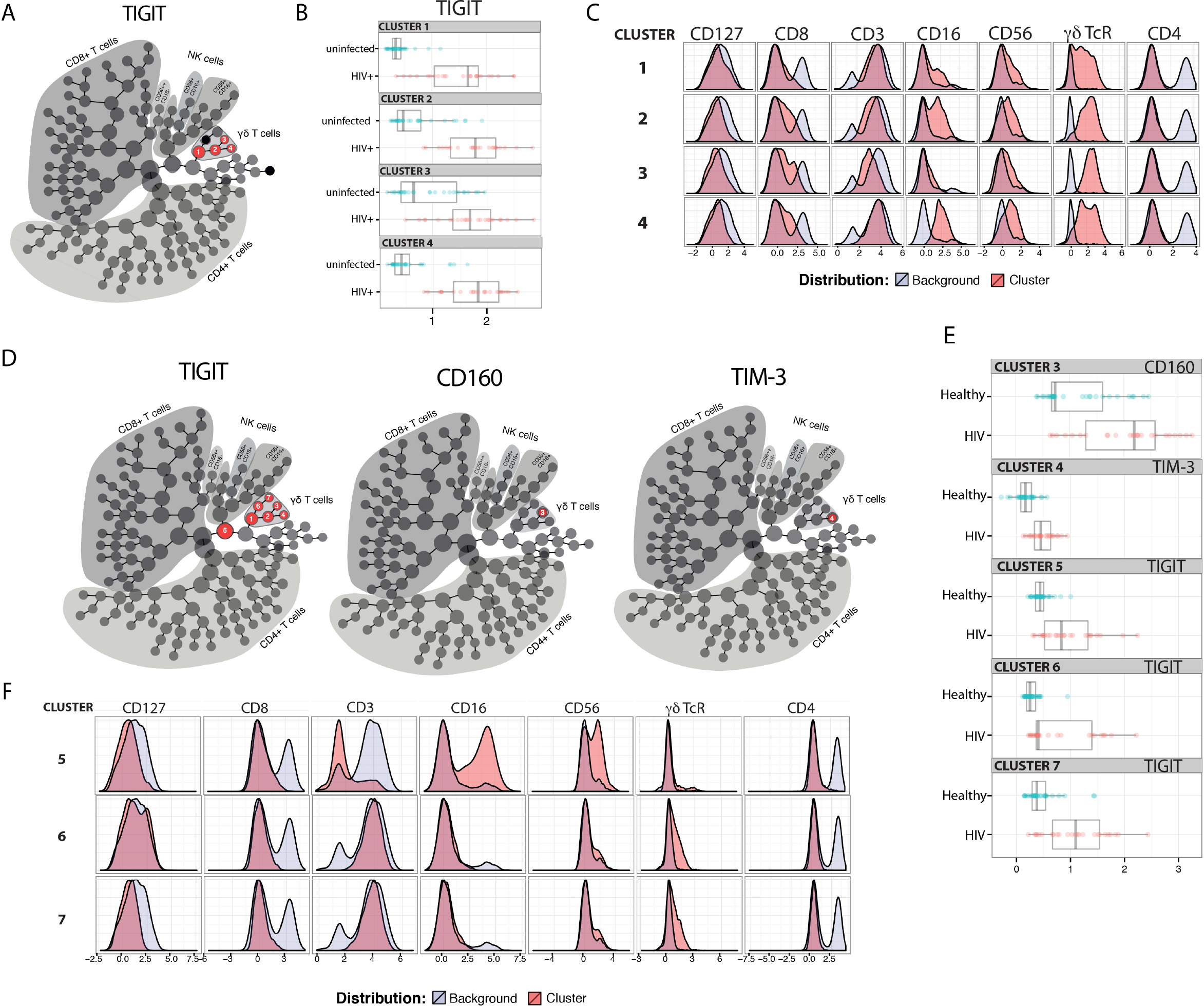
TIGIT, CD160, and TIM-3 on γδ T cells distinguish ART-suppressed HIV+ subjects from uninfected controls. CITRUS analysis of a 16-parameter flow cytometry dataset from PBMC of ART-suppressed HIV+ subjects and uninfected controls. (A) Model selected by minimum cross-validated error rate yielded four clusters necessary to differentiate the groups, numbered 1-4 and highlighted in red. (B) TIGIT expression of cells in clusters 1-4 in the ART-suppressed HIV+ (red) and uninfected control (blue) groups, log-transformed Mean Fluorescence Intensity (MFI) is shown, and each dot represents one subject, (C) expression of other measured antigens on cells from clusters 1-4 as compared to all other (background) cells; (D) Model constrained to include all significant clusters below a FDR threshold of 1% reveals seven clusters with significantly different expression of TIGIT, CD160, and/or TIM-3 between the groups, (E) log-transformed MFI of CD160, TIM-3, and TIGIT expression per subject for clusters 3, 4, and 5-7, respectively, and (F) expression of other measured antigens on cells from clusters 5, 6, and 7. All scales in B, C, E, and F are log-transformed. CITRUS clustering data per lineage channel are shown in Supplemental Fig. 1.

**Table 1.**

| Cohort Characteristics | Uninfected Controls |  | ART-suppressed HIV+<sup>A</sup> |  |
| --- | --- | --- | --- | --- |
|  | Younger (≤35) | Older (≥50) | Younger (≤35) | Older (≥50) |
| n | 21 | 21 | 22 | 28 |
| Age (mean, range) | 27 (22-35) | 58 (50-72) | 28 (22-33) | 58 (50-76) |
| Sex (% M/F) | 100/0 | 100/0 | 100/0 | 86/14 |
| Ethnicity (C/B/A/O)<sup>B</sup> | 15/3/3/0 | 11/6/3/2 | 17/3/1/0 | 18/10/0/0 |
| CD4 count (cells/μm<sup>3</sup>, mean, range)<sup>C</sup> | – | – | 611 (89-1362) | 719 (246-1216) |
| ART duration (years) |  |  | mean n | mean n |
| 0-5 | – | – | 2.2 19 | 2.9 17 |
| 5-10 | – | – | 6.2 3 | 6.8 6 |
| >10 | – | – | – 0 | 16.5 5 |
| Nadir CD4 count (cells/μm<sup>3</sup>) |  |  | mean n | mean n |
| 0-250 | – | – | 159 6 | 133 9 |
| 251-500 | – | – | 339 11 | 311 12 |
| >501 | – | – | 649 5 | 680 6 |

<sup>A</sup>Undetectable HIV-1 viral loads for ≥6 months<sup>B</sup>Caucasian/Black/Asian/other<sup>C</sup>Values within one year of enrollment

γδ T cell IR expression increases with age and HIV infection. We next investigated γδ T cells in more detail by performing traditional expert-driven gating and analysis. We first determined the frequencies and absolute counts of γδ T cells (gating strategy shown in Supplemental Fig. 2A) and found overall similar γδ T cell levels within the HIV+ and uninfected control groups (Supplemental Figure 2B-E). We next measured Vδ1+ and Vδ2+ γδ T cell frequencies from a subset of samples and found an inversion of the Vδ1:Vδ2 ratio in the HIV+ subjects compared to the uninfected group, confirming that the findings in our cohort are consistent with previous reports [45, 62] (Supplemental Fig. 2F-I). Next, we compared the percentages of IR+ γδ T cells between the ART-suppressed HIV+ and control groups and found that the HIV+ subjects expressed significantly higher frequencies of TIGIT+, CD160+, and TIM-3+ cells than age-matched controls (Fig. 2A). As TIM-3 and LAG-3 are not bimodally expressed, setting flow cytometry gates to determine the percentage of positive cells is difficult; therefore, we also calculated the median expression of each IR and found significantly higher levels of TIM-3 but not LAG-3 on the γδ T cells from the HIV+ subjects compared with controls (Fig. 2B). As expression of more than one IR on individual γδ T cells could reflect distinct states of activation or an advanced stage of immune exhaustion [63], we compared the expression of ≥two, ≥three, or ≥four IRs per γδ T cell between our HIV+ and uninfected control subjects and found significantly higher levels in the HIV+ subjects for all comparisons (Fig. 2C). To control for bias due to manual gating, γδ T cell events were extracted from the flow cytometry data using SPADE clustering [64] and γδ T cell IR median expression was assessed (LAG-3 not shown due to negligible results), confirming our manual gating findings (data not shown) and allowing visualization of IR expression from individual subjects (Fig. 2D). Overall patterns indicate higher IR expression of the ART-suppressed HIV+ subjects compared with controls and also seemingly independent regulation of each of the four IRs within individual subjects. Next, subjects were further stratified into the younger and older groups as defined in Table 1 and the percentages of IR+ γδ T cells were compared (Fig. 2E). TIGIT+ γδ T cells were significantly higher in: (1) older versus younger uninfected subjects, (2) HIV+ younger versus uninfected younger subjects, (3) HIV+ older versus uninfected older subjects, and (4) HIV older versus uninfected younger subjects (Fig. 2E). Analysis of the other IRs did not result in significant differences between the groups, with the exception of TIM-3 expression of the uninfected younger versus HIV+ younger subjects (Fig. 2E). Comparison of multi-IR expression with subjects stratified by both age and HIV status shows higher percentages of ≥2IR per cell with advancing age and with HIV infection. Notably, HIV infection and not aging appears to drive γδ T cells to express ≥3 or ≥4 IR per cell (Fig. 2F). Taken together, these results suggest that healthy aging and HIV infection independently drive TIGIT and multi-IR expression on γδ T cells, with HIV exclusively driving γδ T cells to a ≥3 or ≥4IR per cell phenotype. Interestingly, no significant differences in both individual or multi-IR expression were found between the HIV+ young and HIV+ older groups, indicating a lack of a definitive additive or multiplicative effect of both age and HIV on γδ T cell IR expression.

**Figure 2.**
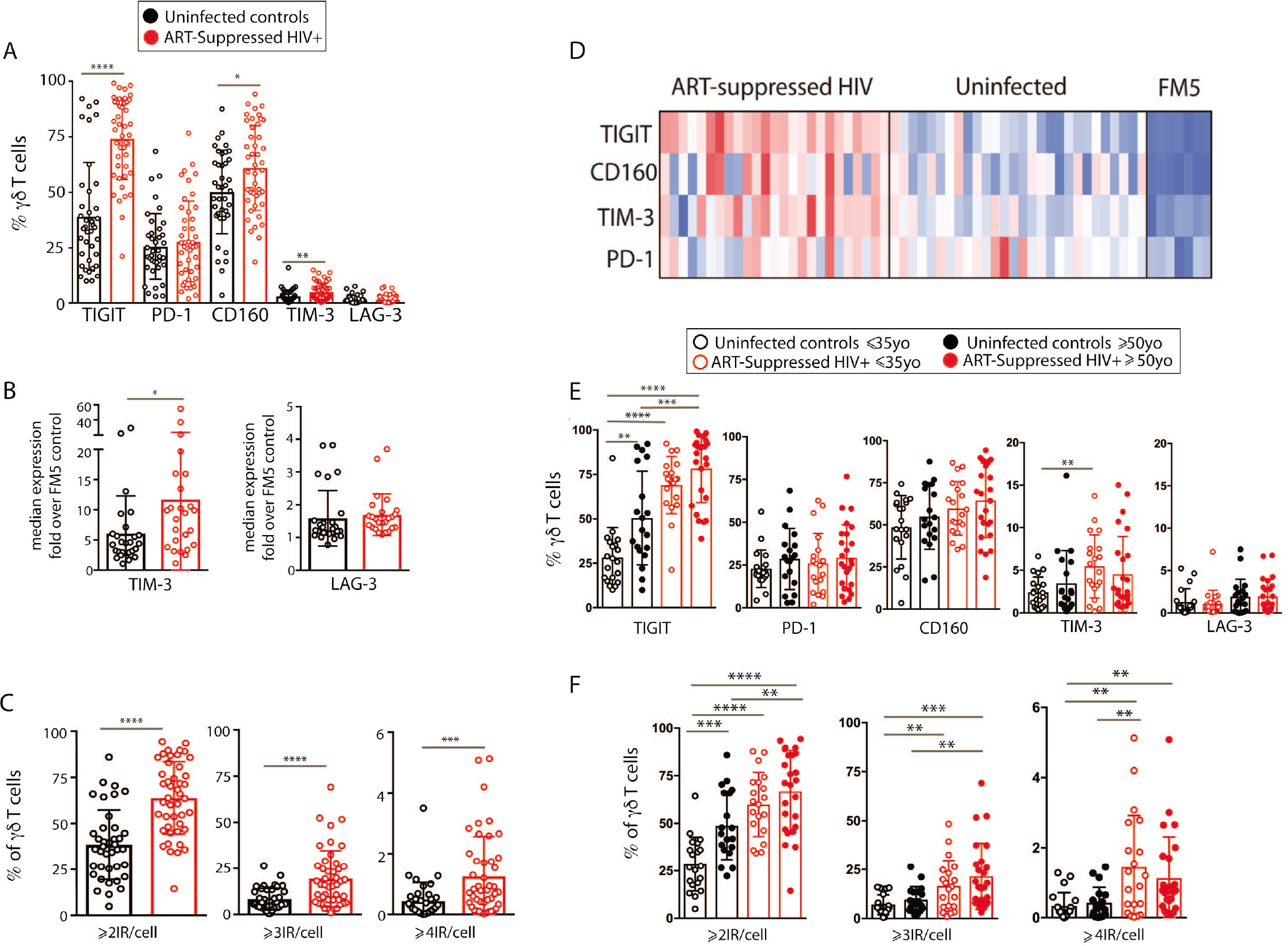
IR expression on γδ T cells from ART-suppressed HIV+ subjects and uninfected controls stratified in young and older groups. (A) The total percentages of IR+ γδ T cells in ART-suppressed HIV+ subjects and control groups; (B) the average fluorescence intensity of γδ T cell TIM-3 and LAG-3 for HIV+ and uninfected subjects, determined by median intensity divided by FM5 control values; (C) the percentage of γδ T cells expressing ≥2, ≥3, or ≥4 IR per cell in HIV+ subjects and controls, (D) median intensity of IR expression on SPADE-identified γδ T cells per individual (one subject per column), including FM5 control samples from each batch run; (E) total percentages of IR+ γδ T cells and (F) percentages of γδ T cells expressing ≥2, ≥3, or ≥4 IR per cell with subjects stratified by both HIV status and age. For A, B, and C, two-tailed t-tests were performed for each comparison. For graphs with multiple comparisons (E, F) only significant results after Bonferroni correction (p<.008)are shown.

γδ T cells expressing distinct combinations of PD-1, TIGIT, and CD160 vary with aging and aviremic HIV infection. We next investigated how the combinational expression of PD-1, TIGIT, and CD160 on γδ T cells differed with healthy aging and ART-suppressed HIV infection. First, the percentage of γδ T cells expressing each of the eight possible combinations of these three IRs were compared across the four subject groups using beta regression of abundance on HIV status and age and correcting for multiple hypotheses testing with a family-wise error rate of 0.05. Both aging and HIV infection were significantly associated with lower percentages of PD-1-TIGIT-CD160-(‘triple negative’) and PD-1-TIGIT-CD160+ (‘CD160 only’) cells. HIV infection was also associated with a lower percentage of PD-1+ TIGIT-CD160-(‘PD-1 only’) cells and a higher percentage of three populations: PD-1-TIGIT+ CD160-(‘TIGIT+ only’), PD1-, TIGIT+ CD160+ (‘TIGIT and CD160 double positive’) and PD-1+ TIGIT+ CD160+ (‘triple positive’) cells (Fig. 3A). Similar to the results for HIV infection, there were higher average frequencies of TIGIT only and TIGIT and CD160 double positive cells in older compared with younger uninfected subjects (albeit not statistically significant). To further assess the potential relationships between γδ T cells bearing different IR combinations, Pearson correlation coefficients were generated for the percentages of all eight possible IR combinations with one another in a pair-wise manner separately for the uninfected controls (Fig. 3B) and ART-suppressed HIV+ subjects (Fig. 3C). Within the uninfected group, strong inverse correlations were found that agree with the results in Fig. 3A, specifically, the triple negative and CD160 only populations with the TIGIT single positive and the TIGIT CD160 double positive cells (Fig. 3B, asterisks). Within the HIV+ group, there were also strong inverse correlations of the triple negative/CD160 only populations, in this case with the TIGIT CD160 double positive cells as well as the triple positive cells (Fig. 3C, asterisks). These data suggest that both aging and HIV infection skew the circulating γδ T cell compartment from predominantly triple negative and CD160 only expressing cells to TIGIT only and CD160 TIGIT double positive cells (Fig. 3D). The triple positive γδ T cell population, while inversely associated with no IR expressing cells in both uninfected controls and HIV+ subjects, is higher in frequency only of the HIV+ subjects compared with controls, and not between the uninfected younger and older groups. Taken together, these data suggest that the triple negative and CD160 only γδ T cell populations are a resting/precursor population to the subsets expressing TIGIT alone, TIGIT with CD160, and TIGIT, CD160, and PD-1, which are likely in an activated or exhausted state (Fig. 3D).

**Figure 3.**
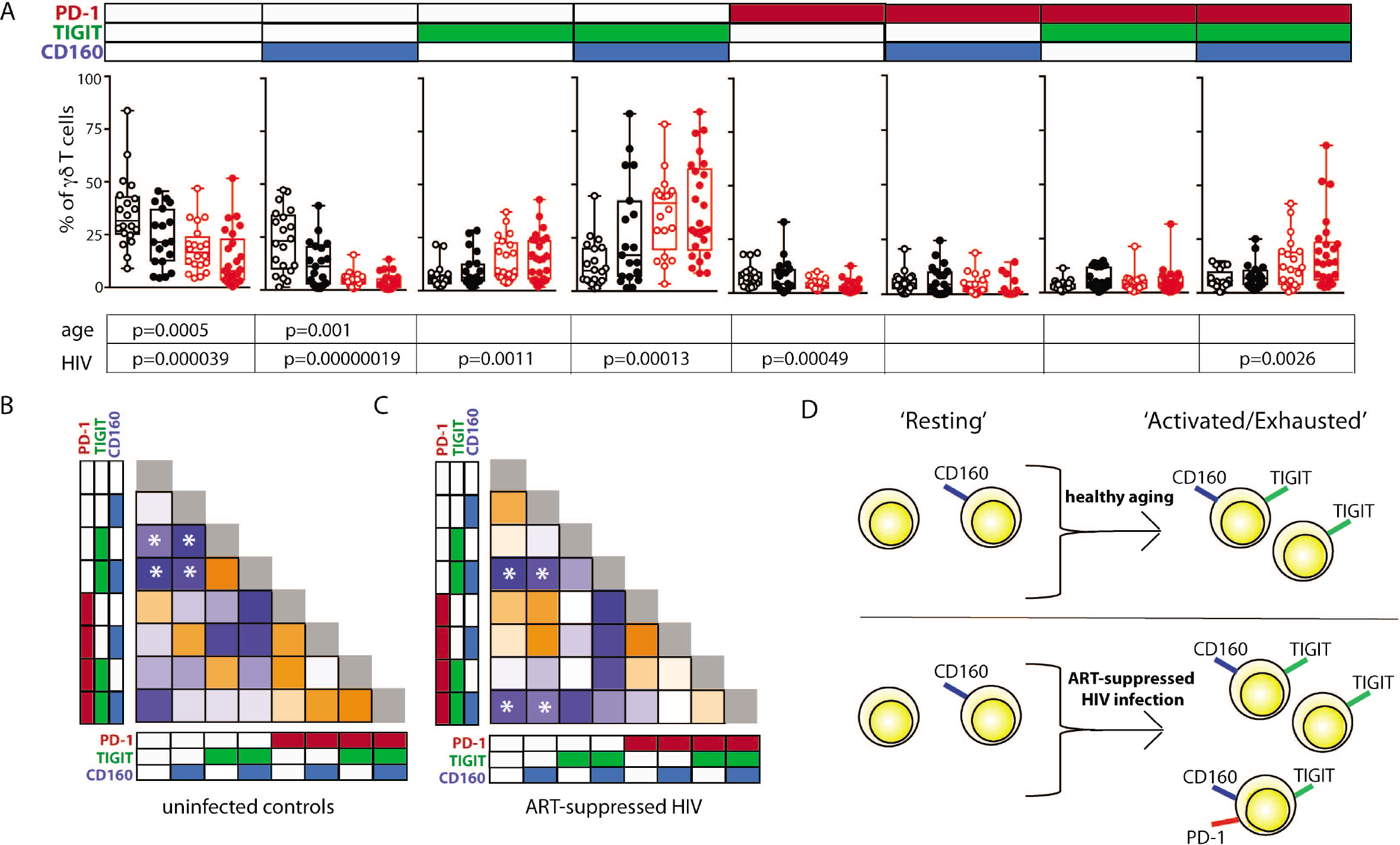
Inhibitory receptor signatures of γδ T cells vary with aging, ART-suppressed HIV infection. (A) gd T cell IR signature analysis of TIGIT, CD160, and PD-1 comparing younger and older subjects and ART-suppressed HIV+ and matched control groups. Abundance for each subset of γδ T cells expressing specific IRs was compared across the four groups using beta regression of abundance on HIV status and age group. Significance of age and HIV infection was determined after correcting for multiple hypotheses testing as described in the Methods section. Due to the low frequencies of γδ T cells positive for either TIM-3 or LAG-3, subsets defined by these IRs were not analyzed separately for this analysis (B) Correlation analysis of the abundances of each IR subset of γδ T cells in uninfected controls and (C) ART-suppressed HIV+ subjects. Heatmaps are colored based on strength of the Pearson correlation coefficient between each IR subset (orange: positive, blue: negative). Both age groups are included in each heatmap. Asterisks indicate biologically interesting correlations used to inform (D) diagram depicting the hypothesized differential progression of IR expression due to healthy aging versus HIV infection.

Spontaneous secretion of inflammatory cytokines and cytotoxic mediators from γδ T cells differentially tracks with TIGIT expression during aging +/-aviremic HIV infection. As an indirect measurement of the potential *in vivo* functional activity of the γδ T cells, cells were sorted and cultured overnight without stimulation and the secretion of 33 analytes was measured in the supernatants. The spontaneous release of sCD137, a marker of cell activation that correlates with circulating C-reactive protein (CRP) in plasma [65] and is positively associated with both acute coronary syndromes [65] and stroke [66], was significantly higher from the γδ T cells of HIV+ older subjects compared with the other three groups (Fig. 4A, results from the other detected analytes are shown in Supplemental Fig. 3). Also, 10 analytes significantly correlated with percentages of defined IR-expressing γδ T cell subsets (Fig. 3D) from either the HIV+ and/or the uninfected subject group (Fig. 4B). Of the HIV+ data, 14 of the 15 significant results included negative correlations with ‘resting’ and positive correlations with ‘activated/exhausted’ populations; this suggests progression from a resting to an activated state with HIV infection. Next, we used simple linear regression to relate levels of eleven analytes detected in the supernatants of a substantial percentage (≥45%) of the subjects with the percentage of TIGIT+ γδ T cells in the paired PBMC sample (eight are shown in Fig. 4C, three in Supplemental Fig. 3). Overall, of γδ T cells from the uninfected controls, the TIGIT+ percentages did not associate with cytokine release (Fig. 4B, C), with the exception of MIP1-β, which showed a negative correlation. Conversely, the percentage of TIGIT+ γδ T cells from the ART-suppressed HIV+ subjects positively and significantly correlated with the secretion of five analytes: sCD137, Granzyme A, Granzyme B, MIP1-β, and CCL20/MIP3-α and showed a positive trend for others such as perforin, TNF-α, and IFN-γ (Fig. 4B, C). Also, there is a notable trend of lower analyte release of γδ T cells from the older compared with younger uninfected subjects, and a converse trend of increased analyte secretion with older age within the HIV+ subject group. These data suggest unlike the uninfected controls, the γδ T cells from ART-suppressed HIV+ subjects are activated, secreting inflammatory factors *in vivo*, and possibly contributing to the inflammatory milieu linked to SNAEs in this population, and that TIGIT expression marks inflammatory activity of γδ T cells from aviremic HIV+ subjects. To further explore the relationships between TIGIT expression and *ex vivo* analyte production by γδ T cells in the HIV+ subjects versus controls, two partial least square regression (PLSR) models (one for uninfected controls, one for HIV+ subjects) were generated to specifically ask if cytokine measurements in the supernatants can predict the percentage of TIGIT+ cells in the PBMC samples. In PLSR modeling, linear combinations of analytes are used to predict the variance in the dependent variables [67, 68], in this case the percentage of TIGIT+ cells. PLSR analysis is more potent than simple regression analyses when the behavior of multiple analytes of interest is known to be interdependent, since it enables determination of the combinations of analytes that maximally correlate with TIGIT expression. Analyte measurements were compressed in low-dimensional latent variable (LV) space; notably, the younger and older subjects separated into distinct groups for each model without the models being trained on age parameter (Fig. 4D, E left panels). Both models significantly linked spontaneous analyte production with TIGIT expression by γδ T cells. VIP (variable importance of projection) calculation was then used to assess the importance of each analyte variable with values greater than one considered influential above average in the model. Notably, the VIP (statistically significant) analytes within each model were different (Fig. 4D, E right panels, light blue bars), confirming that there are distinct associations between TIGIT expression and the cytokines produced from γδ T cells of both HIV+ subjects and uninfected controls. These findings indicate that relationships between γδ T cell TIGIT expression and spontaneous cytokine release change with both normal aging and aging with aviremic HIV infection.

**Figure 4.**
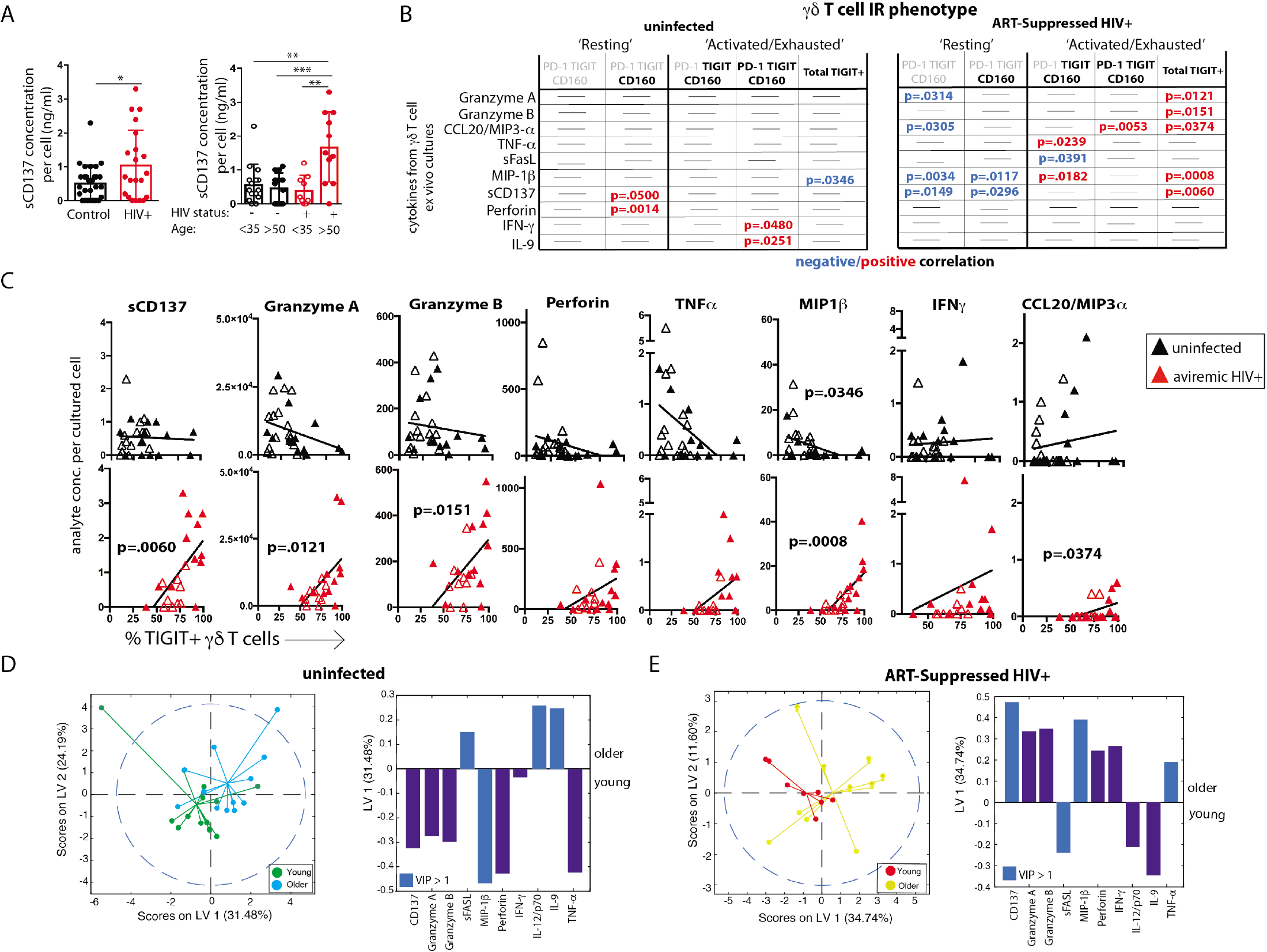
γδ T cell ex vivo spontaneous cytokine secretion profiles reveal differential associations with IR expression and age in ART-suppressed HIV infection and uninfected controls. (A) sCD137 (pg) secreted per cultured γδ T cell from all HIV+ subjects compared with uninfected controls, and with data also stratified into younger and older groups. (B) Tables showing the linear regression analysis results for the 10 analytes that significantly correlated with an IR γδ T cell subset in either subject group (C) Linear regression plots of the percentage of TIGIT+ γδ T cells and cytokine concentration (average per cell) for uninfected controls and ART-suppressed HIV+ subjects for the 8 analytes that highlight the opposite trends between the subject groups (results from the other 3 analytes with >45% subject cells responding is shown in Supp. Fig. 3). Subjects within the younger and older groups are noted with open and solid triangles, respectively. Scores plot and LV1 derived from a PLSR model of supernatant cytokines regressed against the percentage of TIGIT+ γδ T cells for each donor for uninfected controls (D) and ART-suppressed HIV+ subjects (E). Dotted line shows the 95% confidence interval. Units for secreted analytes are listed in Supplemental Figure 3 legend.

Plasma markers of inflammation and coagulation differentially track with TIGIT expression during aging with and without ART-suppressed HIV infection. We measured the concentrations of 16 analytes in plasma, many of which are well defined markers of inflammation and have been strongly associated with the onset of co-morbid conditions and/or mortality in HIV-infected populations [10, 13, 69-72] as well as with disease in normal aging [15-21]. We found significantly higher levels of sCD14, alpha-2 macroglobulin (A2M), fibrinogen, serum amyloid P (SAP), adipsin, and von Willebrand factor (vWF) in the plasma from HIV+ subjects compared to uninfected controls (Supplemental Fig. 4A). When the subjects were further stratified by age, we found significant differences among subject groups for D-dimer, C-reactive protein (CRP), fibrinogen, SAP, adipsin, and vWF (Supplemental Fig. 4B). To determine if there are associations between IR expressing γδ T cell subsets and plasma marker levels, we performed linear regression analysis of the plasma marker data against the percentages of our ‘resting’ or ‘activated/exhausted’ γδ subsets (as defined in Fig. 3D). Of the 16 analytes measured, nine were significantly associated with one or more γδ T cell IR-defined subset(s) in the uninfected controls and/or the ART-suppressed HIV+ groups (Fig. 5A). For both the HIV+ and uninfected subject groups, there was a predominant trend of negative association with the ‘resting’ and positive association with the ‘activated/exhausted’ γδ T cell populations with plasma markers (Fig. 5A-E). Also, it is notable that the younger and older age groups are clearly separated in these analyses among the controls but not the HIV-infected subjects (Fig. 5B-E). These results indicate that TIGIT and multi-IR expression on γδ T cells is linked with the elevated general inflammation found in both healthy aging and ART-suppressed HIV infection, and that age plays distinct roles in these associations with and without ART-suppressed HIV infection. Next, we asked how the concentrations of plasma markers were connected to the percentage of TIGIT+ γδ T cells by training two orthogonalized PLSR models using plasma marker levels to predict the percentage of TIGIT positive cells in the uninfected and HIV+ subject groups (Fig. 5F-G). We note that the models have differing contributions attributed to each marker (as determined by loadings on LV1), and that different markers emerged as the most important to the correlation of inflammatory state to percent TIGIT positive cells (as determined by VIP score); together, these results indicate that the connections between TIGIT+ γδ T cells and the immune network driving systemic inflammation are fundamentally different between aviremic HIV infection and healthy aging. Also, similar to the results in Fig. 4D and E, subjects in the younger and older sub-groups for both uninfected and HIV+ subjects separated without age being added to either model. Overall, our data show that γδ T cells are integral components of distinct inflammatory processes that occur during aging both with and without ART-suppressed HIV infection.

**Figure 5.**
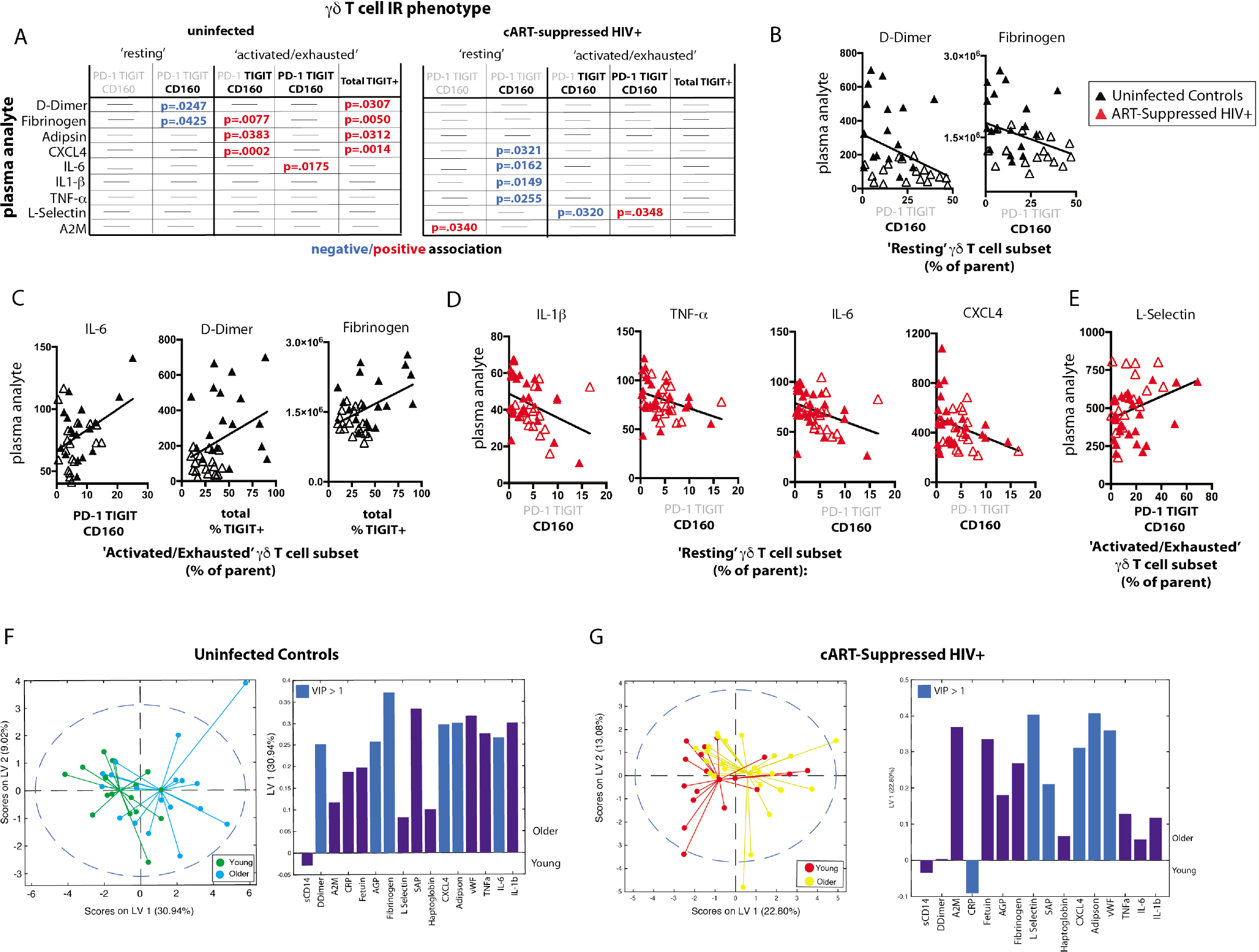
γδ T cell IR signatures correlate with inflammatory plasma markers in both healthy aging and ART-suppressed HIV infection. 16 plasma analytes were measured and linear regression analysis was performed with percentages of IR expressing γδ T cell subsets defined in Figure 3D and total percent TIGIT+; all statistically significant results are shown for uninfected controls and ART-suppressed HIV+ subjects; examples of correlation plots with a fitted linear regression line comparing ‘Resting’ or ‘Activated/Exhausted’ γδ T cell subsets from uninfected controls (black triangles, (A) and (B)) or ART-suppressed HIV+ subjects (red triangles, (C) and (D)) with inflammatory/coagulation markers. Subjects within the younger and older groups are noted with open and solid triangles, respectively, in (B-E). Scores plot and loadings on LV1 derived from a PLSR model of plasma markers versus the percentage of TIGIT+ γδ T cells for uninfected controls (D) and ART-suppressed HIV+ subjects (E). Units for plasma markers are listed in S4 Figure legend.

Partial Least Squares Determinant Analysis (PLS-DA) reveals that γδ IR signatures selectively distinguish the younger and older subject groups within both the uninfected control and ART suppressed HIV+ subject groups. To further confirm that the aging process is distinct in uninfected controls and aviremic HIV+ subjects, we used the percentages of γδ T cells expressing all possible combinations of the IRs PD-1, CD160, TIGIT, and TIM-3 and the 16 plasma marker datasets to train a PLS-DA model that determined if a combination of these variables can separate subjects into four groups based on age and HIV status. In the input data, PLS-DA finds which measurements from each subject would ‘fit’ that subject into its clinical group and verifies if such classification is statistically significant. The chosen biological measurements were sufficient to significantly differentiate all four groups in our dataset (Fig. 6A). In these models, LV1 separated mostly based on HIV status, while LV2 captured the remaining variance attributed mainly to age group. Notably, we trained an alternative model with CD8+ T cell IR signatures used in place of the γδ T cell data and the groups did not separate with significance (data not shown). This analysis indicates that γδ T cells are integral to the inflammatory networks underlying the divergent processes of healthy aging and aging with ART-suppressed HIV infection and we propose that IR expression marks different functional and/or activation/exhaustion states of the γδ T cells in these conditions.

**Figure 6.**
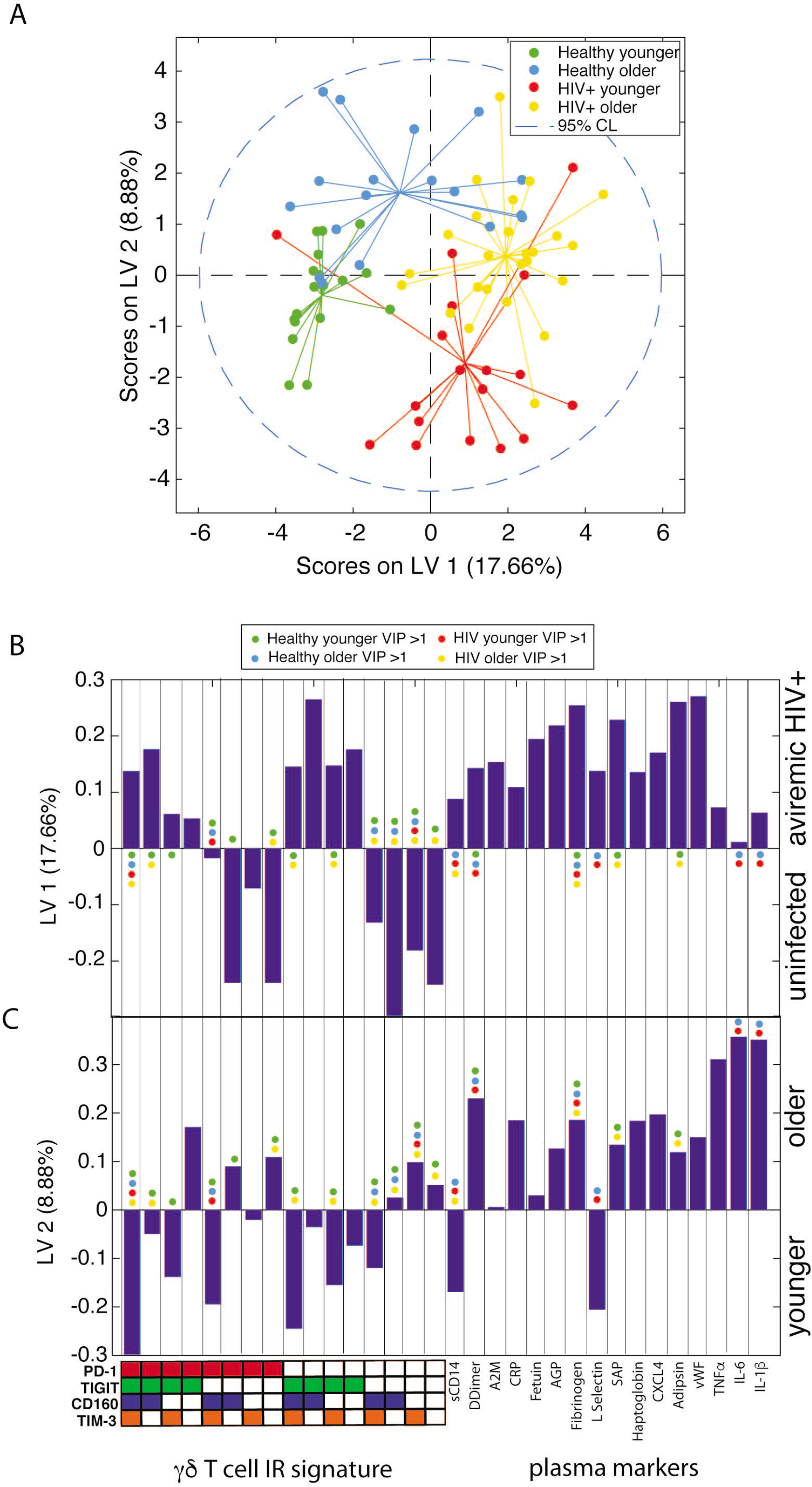
γδ T cell IR signatures and plasma inflammatory cytokines define the divergent aging processes in ART-suppressed HIV+ individuals and uninfected controls. (A) A two-dimensional PLS-DA model constructed using the percentages of all possible combinations of the IRs PD-1, TIGIT, CD160, and TIM-3 and plasma cytokine measurements. Each data point represents scores generated by the model, composed of all measurements for a given patient mapped onto the two-dimensional latent variable space. The percentages on the axes show the percent variance in the dataset captured by a particular LV. Dotted line shows the 95% confidence interval. (B) Bar plot showing the loadings on LV1, the latent variable that separates the scores predominantly based on presence or absence of HIV infection. The Y-axis quantifies the positive or negative contribution of a particular signal to the indicated LV. Colored dots in B, C mark cell populations and cytokines with a VIP score greater than 1 for each subject group in the model. (C) Bar plot as in (B), showing loadings on LV2, the latent variable that separates the scores affected by patient age.

## Discussion

In this study, we present evidence that suggests γδ T cells are an integral component of the inflammatory cycle that leads to non-AIDS morbidities and mortality in aviremic HIV+ individuals, and also, that a distinct inflammatory course is present during aging with aviremic HIV and that is potentially driven by this unique T cell population. Currently, more than 50% of the HIV-infected population in the U.S. is greater than 50 years old [73] and the world population over the ages of 65 and 80 is predicted to double and nearly quadruple, respectively, by 2050 [74]. Elucidating the cell populations and precise immune networks that drive ‘inflamm-aging’ both with and without HIV infection is a preeminent global health priority.

There are multiple proposed triggers of the aberrant inflammation found in HIV+ persons with successful viral suppression, including the latent HIV viral reservoir itself [75], co-infections such as cytomegalovirus (CMV) [76], and translocation of microbial products across the epithelial barrier of the gastrointestinal tract into the systemic circulation [77]. There is a reported connection between harboring latent HIV and IR expression on T cells: CD4+ T cells from virally suppressed HIV+ subjects that express at least one of the IRs TIGIT, PD-1, or LAG-3 contained the majority, on average, of CD4+ T cells with inducible HIV genomes, with multi-IR+ CD4+ T cells particularly enriched for integrated HIV DNA [78]; also, TIGIT+ CD4+ T cell frequencies positively correlate with CD4+ T cell HIV DNA content [49], and TIGIT transcription is elevated in cells bearing replication competent latent HIV [79]. Our recent studies indicate that sensing of HIV viral intron-containing RNA by infected macrophages leads to type I interferon secretion and induction of IRs on co-cultured T cells [80]. Circulating γδ T cells are known to harbor latent virus at a high frequency [47], therefore it is worth investigating if IR+ γδ T cells, particularly TIGIT+ and multi-IR+ subsets, are selectively harboring inducible HIV genomes. Further studies investigating the mechanistic relationship(s) between intracellular HIV, γδ T cell IR expression, and the secretion of inflammatory cytokines and cytotoxic mediators could lead to novel targets for selective therapeutic γδ T cell ablation to help purge the latent HIV pool and reduce general inflammation.

The integrity of the gastrointestinal epithelial barrier is compromised in HIV and SIV infection [77, 81] and does not sufficiently recover with effective ART [82]. This ‘gut leakiness’ is connected to the movement of microbial products such as LPS into the systemic circulation and this likely contributes to general inflammation [83]. γδ T cells are a predominant immune cell subset in intestinal epithelia [24, 25]. Also, the majority of human mucosal intraepithelial lymphocytes (IELs) are Vδ1+ T cells [84, 85], the subset found to predominate the circulation of aviremic HIV+ subjects but not uninfected controls in this study (S2 G-I Fig) and others [45, 62]. *In vivo*, murine Vδ1+ intra-epidermal γδ T cells are activated solely in response to up-regulation of Rae-1, a non-classical MHC stress antigen that binds NKG2D [86]. Also, murine intestinal γδ T cells described as ‘activated yet resting’ expressed very high levels of mRNA for Granzymes A and B [87]. Taken together with our results, these findings lead us to hypothesize that TIGIT expression on circulating γδ T cells of HIV+ subjects may mark cells that have emigrated from the GI tract to the systemic circulation and that this normally tissue resident Vδ1+ subset, activated and armed with cytolytic machinery, contributes to general inflammation. Also, activation or direct infection of tissue resident γδ T cells by HIV virions within the gut-associated lymphoid tissue (GALT) could lead to the loss of epithelial barrier integrity and/or cause other mucosal aberrancies that promote systemic inflammation. Immune cell composition in the blood has reflected what is found in the GALT in aviremic HIV+ subjects [88]. Future work examining mucosal resident γδ T cells in SIV-infected animals and HIV+ individuals could provide new insight into how immune changes in the gut direct general inflammation and SNAEs in aviremic HIV infection.

An immune cell subset strongly linked to inflammation and SNAEs in aviremic HIV+ infection is monocytes/macrophages. Markers of monocyte/macrophage activation in the circulation, such as sCD14, sCD163, and tissue factor (TF), have been linked with mortality, atherosclerosis, and other markers of inflammation and coagulation [89-91]. Furthermore, TF-expressing monocytes, present with successful HIV viral suppression, produce multiple inflammatory cytokines upon exposure to LPS and induce a coagulation cascade. Treatment of SIV-infected macaques with an anti-coagulant that blocks the TF pathway led to a decrease in circulating D-dimer levels as well as markers of immune activation [92]. Unlike traditional (adaptive) T cells, a principal means of γδ T cells activation is via cytokine and NK receptors [34], mechanisms easily triggered by activated monocyte/macrophage populations. The effect of TF on γδ T cells is unknown to date. It is of interest to investigate the potential cross-activation of monocytes/macrophages and γδ T cells in aviremic HIV infection, as aberrantly activated γδ T cells may induce monocyte/macrophage activation/TF expression, or conversely, inflammatory monocyte/macrophage populations may directly cause the activation of γδ T cells.

Inhibitory receptors, such as PD-1, TIM-3, and TIGIT, negatively regulate T cell activation and are believed to be critical for limiting immunopathology during immune responses. Expression of IRs is often associated with diminished functional capacity, frequently called immune exhaustion, in HIV, aging and other chronic diseases [50, 53, 59]. TIGIT (T cell immuno-receptor with immunoglobulin and ITIM domains) is a more recently described inhibitory receptor that is a member of the CD28 family of signaling molecules [93]. It is expressed on NK cells, CD8+ T cells, and CD4+ T cells [93], is induced upon activation in multiple immune cell subsets [94-96], and its expression marks exhausted CD8+ T cells in HIV and SIV infection [49]. Publications investigating IR expression on γδ T cells are extremely limited to date and include a report of PD-1 expression on γδ^lo^ T cells in mouse skin [97] and another of TIGIT expression on γδ T cells in the epidermis of stem cell transplant recipients [98]. In our study, TIGIT expression tracked with active, *ex vivo* secretion of inflammatory cytokines and cytotoxic mediators, without re-stimulation, from the γδ T cells of aviremic HIV+ subjects. These findings suggest that TIGIT+ γδ T cells are activated in ART-suppressed HIV+ subjects and secrete inflammatory mediators *in vivo*. This association was not found in the uninfected control subjects; on the contrary, the cytokine data from controls indicate that the increased γδ T cell IR expression in healthy aging is linked with diminished cytokine secretion (Fig. 4). However, it is important to note that while the percentage of TIGIT+ γδ T cells sorted into the individual culture wells significantly correlates with the amount of inflammatory analyte measured, the functional profiles of actual TIGIT+ versus TIGIT-γδ T cells were not directly measured in this study. It can be concluded that IR expression by different immune cell subsets in the same anatomical site may reflect contrary functional states, and also, the same immune cell subset with an identical IR signature may have very different functionality in different disease states or other conditions. Therefore, we contend that IR expression analysis must be consistently paired with functional readouts to determine the true status of a given immune cell population. As IRs have been referred to as “exhaustion markers” [99, 100]; our data support arguments by others to not limit IR definitions in this manner [101].

It has been postulated that multi-IR expression by individual T cells reflects more advanced immune exhaustion [102] in agreement with reports of T cells in virally infected mice [63, 103]. In our study, the percentages of γδ T cells expressing ≥2IR per cell were higher with both healthy aging and HIV infection, yet the ≥3IR and ≥4IR per cell levels were elevated only with HIV infection (Fig. 2C, F) possibly suggesting a more progressed immune exhaustion status of these cells. However, our measurements of analyte secretion indicate that the HIV+ subjects’ γδ T cells expressing IRs are activated and not exhausted (Fig. 4B). When we performed more detailed IR signature analysis to parse out potential lineage trajectories of γδ T cell subsets bearing combinations of PD-1, TIGIT, and CD160 in ART-suppressed HIV infection and/or aging, interesting findings emerged; firstly, our results show that a substantial proportion of the circulating γδ T cell compartment in healthy younger subjects express CD160 (Fig. 3A) that is not elevated with HIV- or age-associated activation or exhaustion (Fig. 3A-C). CD160 recognizes class 1a and 1b molecules [104] and serves as a co-receptor for activation of γδ T cells [105] yet its expression has been linked to immune exhaustion on CD8+ T cells in HIV and other chronic diseases [106, 107]. These results underscore the importance of evaluating IR expression in a cell type and disease-specific manner. Also, our data indicate that TIGIT and PD-1 are up-regulated on what we cautiously define as ‘activated/exhausted’ cell subsets, and the PD-1+ TIGIT+ CD160+ (triple positive) T cells are elevated with aviremic HIV infection and not healthy aging, suggesting that HIV drives further progression of γδ T cell populations along activation/exhaustion routes. Taken together, our data suggest that multi-IR expression per cell does not always signify advanced immune exhaustion and that combinational IR signature analysis may provide new insight into the progression of changes in immune cell functional activity and capacity (activation versus exhaustion) in chronic conditions. Future work measuring the functional profiles of sorted γδ T populations with distinct IR signatures from our cohort will help elucidate how functions change with IR expression during aging with and without aviremic HIV infection.

There is debate in the literature as to whether aviremic HIV infection actually accelerates aging (as defined via age-associated inflammation/immune changes and morbidities) [108-110]. If so, we would expect similarities in immune signatures between the younger HIV+ and healthy older subjects, as well as between the HIV+ younger and HIV+ older individuals. Some of our data, specifically the γδ T cell IR signatures compared between the HIV+ younger and HIV+ older groups (Figs. 2, 3), agree with this concept and suggest that aviremic HIV infection and aging are not impacting immune cell phenotypes in an additive or multiplicative manner, consistent with a report of CD8+ T cells [111]. However, when looking beyond surface marker composition, our data indicate key differences between the γδ T cells in all four of our subject groups. For example, γδ T cell secretion of sCD137 is significantly higher from the cells from the HIV older group compared to all other three subject groups (HIV younger, uninfected younger, and uninfected older) (Fig. 4A), indicating a synergistic impact of age and HIV on the activation of γδ T cells. sCD137 is a pro-inflammatory molecule that can be produced as an alternatively spliced variant or by proteolytic cleavage of CD137 (4-1BB) [112], a member of TNF superfamily, and a marker associated with multiple inflammatory diseases. sCD137 plasma levels positively associate with acute coronary syndromes [65] and atherothrombotic stroke [66], and we hypothesize that *in vivo*, γδ T cells produce sCD137 which is also known to trigger pro-inflammatory networks that involve macrophages and DCs [113] Also, our PSLR results show that the relationships between γδ IR signatures and both the cytokines they produce and plasma markers with which they co-vary are different in the younger versus older subjects in both the uninfected control and HIV+ groups, since the younger and older subjects separated in our PLSR models without age being added as a variable. These results show that there are distinct aging courses in relation to immune network changes with and without HIV infection. Finally and perhaps most importantly, in our PLS-DA model all four subject groups separated with significance (Fig. 6) demonstrating separate aging processes with and without HIV infection.

Scientific research is trending away from hypothesis-driven approaches and more toward the systematic collection of large genomic, proteomic, and other datasets. There are notable advantages to larger multi-parameter data collection over more scientifically focused research approaches; for example, the substantive efficiency in the collection of large amounts of data from rare and/or limited samples. A current drawback to more extensive data collection is the analysis bottleneck that exists using most current expert-driven (manual) methods. Here, we demonstrate the utility of multiple bioinformatic algorithms to glean key differences between subject groups that then enable next-step focused biological (mechanistic) exploration, elucidation of the putative cross-impact(s) of co-existing conditions, and potentially improved paths to biomarker discovery. For example, use of CITRUS [61] enabled discovery of the marker (TIGIT) and immune cell subset (γδ T) that could stratify individual subjects into the correct subject group (HIV+ versus control) (Fig. 1). Subsequently, immune signature datasets collected from multiple techniques and biological sources (including flow cytometry of PBMC and multiplex assays of plasma samples) were used in additional algorithms in this study to reveal the relationships among measured variables and various outcomes (PLSR and PLS-DA). Our PLSR and PLS-DA results revealed the differential impact of aging with and without aviremic HIV, but they also highlight the widespread multivariate shift affecting multiple variables in HIV and aging rather than isolated changes in plasma levels of inflammatory markers or single IR expression. Biomarkers, albeit predictive, prognostic, or pharmacodynamic, are difficult to find with the required specificity and sensitivity to lead to patient benefit; alternatively, it has been proposed that using combinatorial biomarker signatures may be more efficacious [114]. Therefore, we propose that the acquisition of large immune signature datasets and addition of such data into multivariate computational algorithms is a potent workflow to enable improved biomarker discovery.

Altogether, our findings indicate that γδ T cells are a possible source of inflammation in aviremic HIV infection and potentially other inflammatory diseases, as well as healthy aging. IR expression on circulating γδ T cells, and TIGIT in particular, should be investigated for biomarker utility to effectively diagnose, predict, and/or treat both general inflammation and propensity to age-associated morbidities and mortality with and without HIV infection. Therapeutics to enhance the Vδ2+ γδ T cell subset have been proposed [115]; alternatively, our data suggest that targeted depletion of TIGIT+ Vδ1+ cells, as it is possible that they are highly inflammatory emigrants from the gastrointestinal system, could be effective at reducing inflammation and improving the quality and lifespan of individuals living with HIV.

## Material and Methods

### Participants

The subjects were part of the HIV and aging cohort which enrolled HIV+ subjects with ART effective viral suppression (undetectable HIV-1 RNA <50 copies/ml) for a minimum of six months and were age-stratified into≤35 years and ≥50 years age groups from the infectious disease clinics at Brigham and Women’s Hospital, Beth Israel Deaconess Hospital, Massachusetts General Hospital, and Boston Medical Center, all located in Boston, MA. Study protocols were approved by the institutional review boards at each institution and all subjects provided written informed consent. The committee that approved the research protocol was the Boston University Institutional Review Board, IRB# H-33095. Uninfected controls were recruited from the same clinics in Boston and had similar demographic and socioeconomic status as HIV+ subjects. HIV negative status was verified with both a negative HIV antibody test and HIV-1 viral load test. Exclusion criteria included active hepatitis B or C, recent active infection within past six months, recent immunomodulatory therapy, and receipt of an HIV vaccine. HIV+ subjects in different age groups are matched by duration of ART (Table 1). Detailed information about the study subject characteristics can be found in Table 1. All subjects were recruited with detailed clinical history and socio-demographic data, and information of other common confounders, such as gender, co-infections, and other pro-inflammatory exposures (i.e. STDs, obesity, and drug/tobacco use).

### Peripheral blood processing

Peripheral blood mononuclear cells (PBMC) were isolated from blood originally drawn into EDTA tubes by Ficoll-Hypaque and cryopreserved in liquid nitrogen until use. Plasma was acquired from whole blood and stored at −80°C until use. All plasma and PBMC samples used in this study are from the same blood draw.

### Flow Cytometry Panels and Reagents

Two flow cytometry panels were used in the study. Panel 1 was a modified version of the previously published panel [116] where 3 markers were moved to different detection channels and one replaced. Panel 2 was a modification of Panel 1 constructed to allow detection of γδ T cell Vδ1 and Vδ2 subsets. All reagents used in both panels are listed in Supplementary Table I. Buffers and blocking reagents were used as described [116]. All fluorescent reagents were titrated and Fluorescence Minus One (FMO) controls were performed to verify the validity of the modified panels. Ultracomp capture beads (Thermo Fisher) were used to establish the compensation matrix. Ultra Rainbow Beads (Spherotech) were used to establish target values for running the instrument. CS&T beads (BD) were used for daily QC of the instruments.

### Flow Cytometry Protocol

1×10^7^ PBMC per donor were thawed and immediately stained with a 16-color antibody/reagent cocktail developed for the BUMC FACSARIA (BD Biosciences) cell sorter as previously described [60]. Briefly, cells were washed, stained with live/dead reagent (Biolegend’s Zombie NIR) in DPBS for 20 min, washed again with FACS buffer (DPBS/0.5% BSA/2mM EDTA), and pre-blocked with human Fc-blocking reagent (Biolegend) for 10 min. The antibody cocktail was prepared by combining antibody stocks with BD Brilliant Buffer and adding the master mixes to the pre-blocked cells that were then incubated for 20 minutes at room temperature. After staining, cells were washed twice with FACS buffer (PBS 0.5% BSA 2mM EDTA), resuspended in FACS buffer and run on the BUMC FACSARIA cell sorter (BD Biosciences). Samples were divided into 11 different runs and a mix of subjects from the four groups (uninfected younger, uninfected older, ART Suppressed HIV+ younger, ART-suppressed HIV+ older) were included in each run to control for potential alterations in machine performance and batch effects. For each run, a mixture containing 5% of each individual sample was prepared and stained with the reagent mix that did not include IR-specific fluorescent antibodies (Fluorescence Minus Five, or FM5). For Panel 2, anti-Vδ2-biotin and anti-Vδ1-PE added were added to cells for 20 minutes then cells were washed prior to the addition of other reagents that were then added and incubated for 20 minutes. For each batch run, Ultracomp capture beads (Thermo Fisher) were stained with same reagents and acquired prior to the samples. PDGFRα-APC antibody (Biolegend) was used for the APC single stain.

γδ T cell cultures: During data acquisition and sample running on the BD FACSARIA, γδ T cells (LIVE/DEAD-, CD14-, CD19-, CD3+, gdTCR+) were sorted into tubes and cultured in 25 ml of RPMI1640 10% FBS / 1% pen-strep in half-area 96-well flat-bottom plates (Greiner) for 20 hours. The number of cells cultured per well varied from 2000 to 26000, with a comparable range in numbers across each of the four subject groups. At the end of the culture period, supernatants were frozen into multiple aliquots for inflammatory and coagulation marker analysis.

### Milliplex Bead Assays of culture supernatants

Supernatant samples were thawed, centrifuged for 30 seconds to remove debris, and applied to a 384-well plate for analyte measurement via use of the Milliplex human Th17 25-plex kit (Millipore Sigma) and a modified Milliplex human CD8 9-plex kit (Millipore Sigma). Antibodies and magnetic beads were diluted 1:1 with assay buffer and utilized at half-volume to adjust the manufacturer’s protocol to the 384-well plate format. Plates were washed in between incubations using a BioTek 406 Touch plate washer (BioTek) and read using the Luminex FlexMap 3D system (Luminex). Observed concentration values were adjusted to cell count.

### Milliplex and ELISA Assays of plasma samples

Frozen plasma samples were thawed and analyzed using (1) the 11-plex Human Cardiovascular Disease (Acute Phase) Milliplex kit (MilliporeSigma, HCVD3MAG-67K), (2) a custom 3-plex Milliplex kit for TNF-α, IL-6 and IL-1β (HCYTOMAG-3K) sCD14 and D-dimer ELISA kits (from RD Systems and Abcam, respectively). For (1), samples were pre-diluted 1:40000 and the kit was run according to manufacturer’s instructions and as described above.

### Flow Cytometry data analysis

Data were recorded with BD FACSDIVA 6.1.2 and automatically compensated with compensation matrix recorded within the sample files according to FCS 3.0 standards. Manual gating (expert-guided analysis) was performed in FlowJo v.10.2-10.3 (FlowJo, Inc) while blinded to sample status. Inhibitory receptor (IR) gating was guided using the FM5 sample from each run described above. Numeric values from FlowJo data analysis were exported to Prism 7.0 (Graphpad) for statistical analysis. Flow cytometry data files were also uploaded to Cytobank, a cloud-based computational platform, and arcsin-transformed. Doublets, dead cells, debris, CD14+ cells and CD19+ cells were excluded by manual gating and remaining live single non-CD14, non-CD19 events (downsampled to 10,000 events/subject) were input into the CITRUS algorithm [61]. After hierarchical clustering based on lineage markers (CD3, CD4, CD8, γδ TCR, CD127, CD16, CD56) shown in Fig. S1, median fluorescence of TIGIT, TIM-3, PD-1, CD160 and LAG-3 expression was measured and a supervised comparison between HIV- and HIV+ individuals was carried out and assessed via Nearest Shrunken Centroid (PAMR) association modeling (Fig S1). Additionally, live, single, non-CD14, non-CD19 events were input to the SPADE clustering algorithm, and the nodes containing cells with higher γδ TCR expression were exported from each sample file and used for IR median fluorescence intensity representation (Figure 2D). Beta regression of abundance on HIV status and age group was performed in Figure 3A. To correct for multiple hypothesis testing, a significance threshold was set for each regression term using a family-wise error rate of 0.05. The significance threshold was established in the following manner. First, 100 permuted abundance datasets were created by randomly resampling without replacement the abundance of each of the eight IR subsets within each subject, so that the inter-dependency of the IR subset abundances for an individual was maintained. Then, for each of the 100 permuted abundances, the beta regression for each IR subset was performed and the minimum p value across all IR subsets was recorded for each model term. Finally, the 5^th^ percentile of this distribution of minimum p values was set as the significance threshold for each model term (HIV threshold: 0.0047; age group threshold: 0.0068).

### PLS analysis

Partial least squares discriminant analysis (PLSDA) and partial least squares regression (PLSR) were performed using the MATLAB PLS Toolbox (Eigenvector Research, Inc.). Data were normalized along each X and Y parameter by Z-score before application of the algorithm. Cross-validation was performed with one-third of the relevant dataset. The number of latent variables (LVs) was chosen so as to minimize cumulative error over all predictions. We orthogonally rotated the models so that maximal separation was achieved across LV1 where noted. We calculated model confidence by randomly permuting Y 100 times and rebuilding the model to form a distribution of error for these randomly generated models. We compared our model to this distribution with the Mann-Whitney U test to determine the significance of our model. The importance of each parameter to the overall model prediction was quantified using variable importance in projection (VIP) score. A VIP score greater than 1 (above average contribution) was considered important for model performance and prediction.

## Author Contributions

AB, AO, NL, and JSC conceived and designed the experiments; KD, DL, NL contributed to reagents, materials, and funding; AB, RP, JSC performed experiments; AB, AS, KD, EP, RP, and JSC analyzed data; and AB and JSC wrote the manuscript. All authors read and approved the final version of the manuscript.

## Data availability

The raw data supporting the conclusions of this manuscript will be made available by the authors, without undue reservation, to any qualified researcher.

## Conflict of interest statement

The authors declare that the research was conducted in the absence of any personal, professional, or financial relationships that could potentially be construed as a conflict of interest.

## Acknowledgements

We thank Drs. Jeffrey Browning, Amedeo Cappione, Andrew Henderson, and Barbara Nikolajczyk for their critical reading of the manuscript, Drs. Suryaram Gummuluru and Manish Sagar for their helpful discussions regarding data interpretation, and Dr. Barbara Nikolajczyk and Dr. Madhur Agrawal for their assistance with the multiplex assays. All flow cytometry data were collected at the Boston University School of Medicine Flow Cytometry Core Facility.

## Funding

This work was supported by the National Institute of Health (R01-DK108056, R01 DA041748-01,1UL1TR001430), the National Institute Of General Medical Sciences (R44GM117914), and the Army Institute for Collaborative Biotechnologies (W911NF-09-0001).

---

**Supplemental Table 1.** Flow Cytometry Reagents

| **Specificity** | **Conjugate** | **Ab clone** | **Vendor** | **Panels** | **Catalog number** |
| --- | --- | --- | --- | --- | --- |
| CD25 | BUV 395 | 2A3 | BD | 1 | 564034 |
| Streptavidin | BUV 395 |  | BD | 2 | 564176 |
| CD127 | BUV 737 | HIL-7R-M21 | BD | 1 | 564300 |
| CD8 | BUV 805 | SK1 | BD | 1, 2 | 564913 |
| PD-1 | BV 421 | EH12.2H7 | Biolegend | 1, 2 | 329919 |
| CD3 | BV 510 | OKT3 | Biolegend | 1, 2 | 317331 |
| CD16 | BV 605 | 3G8 | Biolegend | 1, 2 | 302039 |
| Tim3 | BV 650 | 7D3 | BD | 1, 2 | 565565 |
| CD56 | BV 786 | NCAM16.2 | BD | 1, 2 | 564058 |
| CD160 | AF488 | BY55 | Thermo Fisher | 1, 2 | 53-1609-41 |
| Vα24 TCR | PE | C15 | Beckman Coulter | 1 | COIM2283 |
| Vδ1 TCR | PE | REA173 | Miltenyi | 2 | 130-100-536 |
| LAG3 | PE-e610 | 3DS223H | Thermo Fisher | 1 | 12-2239-41 |
| CD27 | PE-e610 | 0323 | Thermo Fisher | 2 | 61-0279-41 |
| TIGIT | PE-ef710 | MBSA43 | Thermo Fisher | 1, 2 | 46-9500-41 |
| $\gamma\delta$TCR | PE-Cy7 | B1 | Biolegend | 1, 2 | 331221 |
| CD1d tetramer | APC | n/a | NIH tetramer facility | 1 |  |
| Vγ9 TCR | APC | B3 | Biolegend | 2 | 331309 |
| CD4 | AF700 | RPA-T4 | Biolegend | 1, 2 | 300526 |
| CD14 | APC-Cy7 | HCD14 | Biolegend | 1, 2 | 325619 |
| CD19 | APC-Cy7 | HIB19 | Biolegend | 1, 2 | 302217 |
| NIR Zombie | n/a | n/a | Biolegend | 1, 2 | 423105 |
| Vδ2 | biotin | B6 | Biolegend | 2 | 331404 |
| Brilliant Stain Buffer | n/a | n/a | BD | 1, 2 | 563794 |
| Human TruStain FcX FcR Block | n/a | n/a | Biolegend | 1, 2 | 422301 |

**Supplemental Figure 2.**
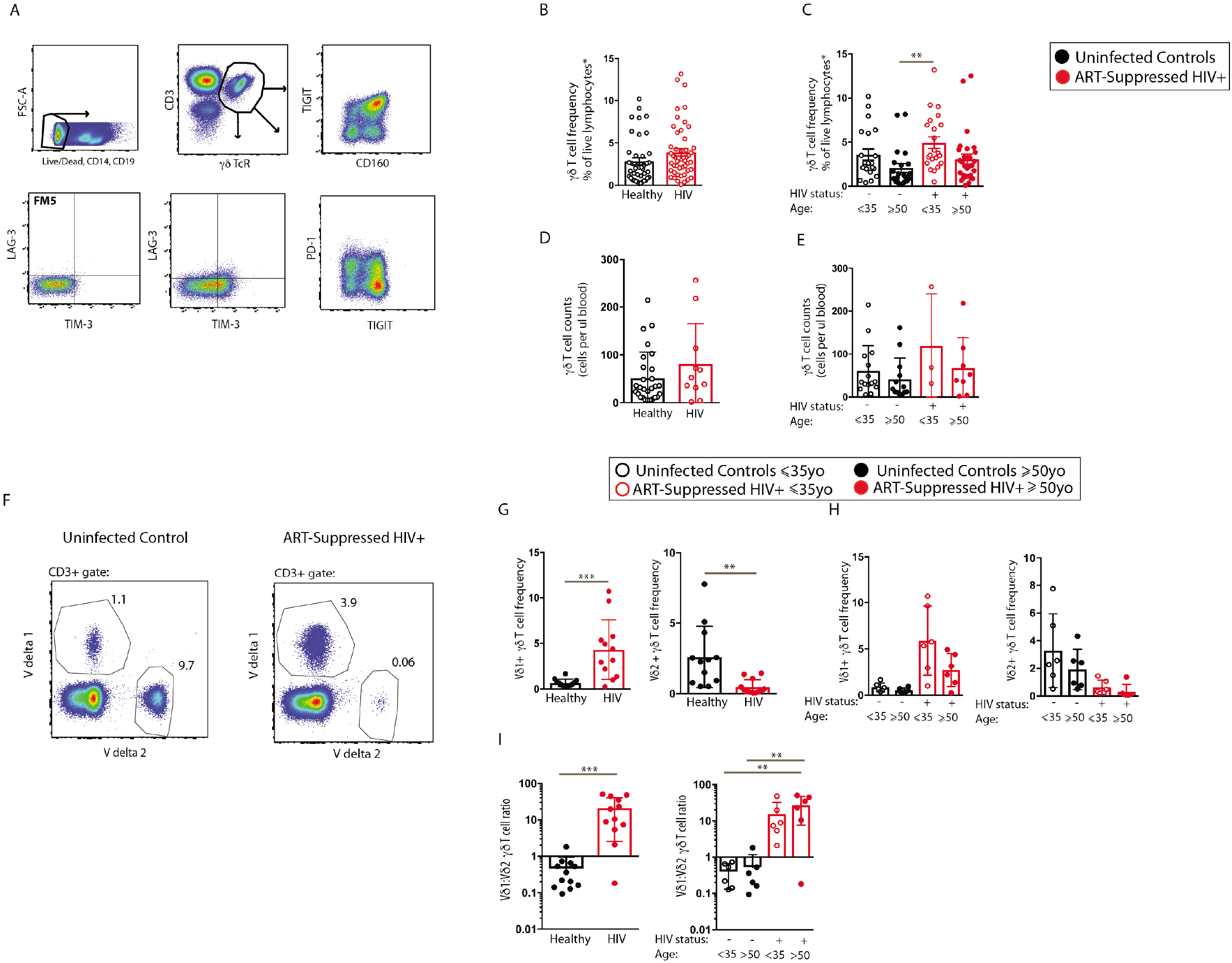
γδT cell gating strategies, frequencies, absolute counts, and Vδl, Vδ2 subsets from subjects of the HIV and Aging cohort. (A) Flow cytometry gating strategy used to enumerate circulating γδT cells and γδT cell inhibitory receptor expression, FM5 (Fluorescence Minus 5) is staining with all reagents in the panel except the antibodies for the five inhibitory receptors. γö T cell frequencies (B) and absolute counts (D) within PBMC from ART-suppressed HIV+ and uninfected control groups. γδT cell frequencies (C) and absolute counts (E) of the HIV+ subjects and uninfected controls further stratified by age. (F) Staining example of γδT cells for Vδ1, and Vδ2 from a HIV+ and uninfected control subject. (G) Vδ1 and Vδ2 frequencies of live lymphocytes from the total ART-suppressed HIV+ and uninfected control groups and (H) from the same subjects when further stratified by age. (I) Vδ1: Vδ2 ratio per subject of the uninfected control and HIV+ subject groups both combined and stratified by age. Two-tailed T tests were performed for statistical analysis. *p<.O5, **p<.01, ***p<.001, ****p<.0001.

**Supplemental Figure 3:**
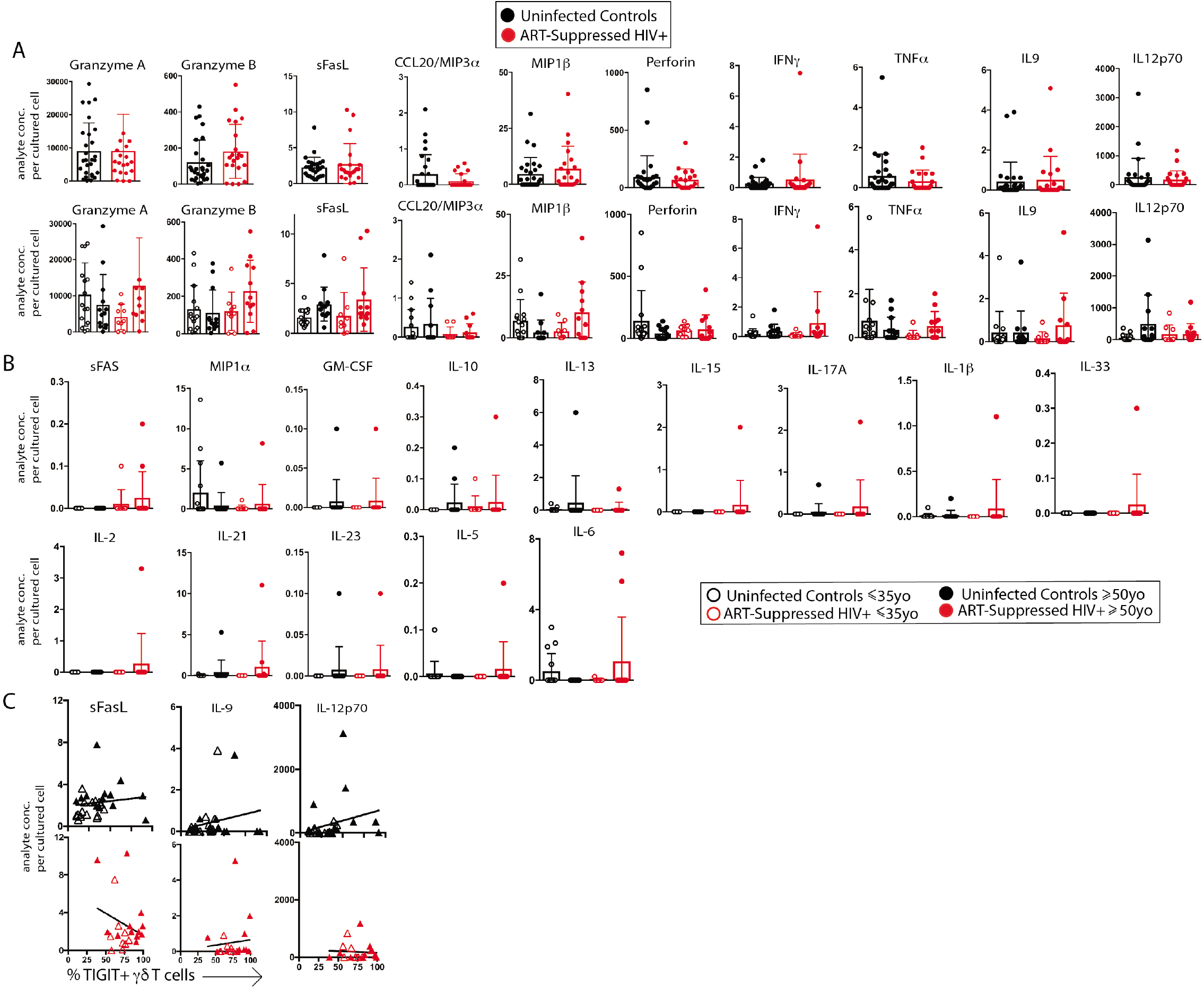
Measurement of spontaneous release of 32 analytes from γδ T cells from the BUMC HIV and Aging cohort: additional data not included in Fig. 4. (A) 11 analytes were detected in the supernatants of >45% of all subjects, including sCDI37 (data shown in Figure 4) and ten others shown in (A). There were no significant differences between the groups. In (B) analytes with detectable levels in at least one subject are shown. (C) analyte concentrations versus % TIGIT+ cells in culture for sFASL, IL-9, and IL-12p7O. Subjects within the younger and older groups are noted with white and solid triangles, respectively. There were no detectable amounts of I LI 7F, IL-22, IL-4, IL-17E/IL-25, IL-27, IL-31, TNF-ß, and IL-28A produced from yδ T cells sorted from any subjects. Units are as follows: sCD137, sFAS, sFASL, Granzyme A, Granzyme B, IL-6, Ml P-1 α, MIP-1 ß; pg/cell; IL-17, IL-4, GM-CSF, IL-27, IL-22, TNF-ß, IL-21; ng x 104 per cell; IFN-y, IL-10, MlP-3α, IL-15, IL-1 ß, IL-33, IL-21, IL-13, IL-17A, IL-2, IL-9, IL-5, IL-6, TNF-α; pg x104 per cell.

**Supplemental Figure 4.**
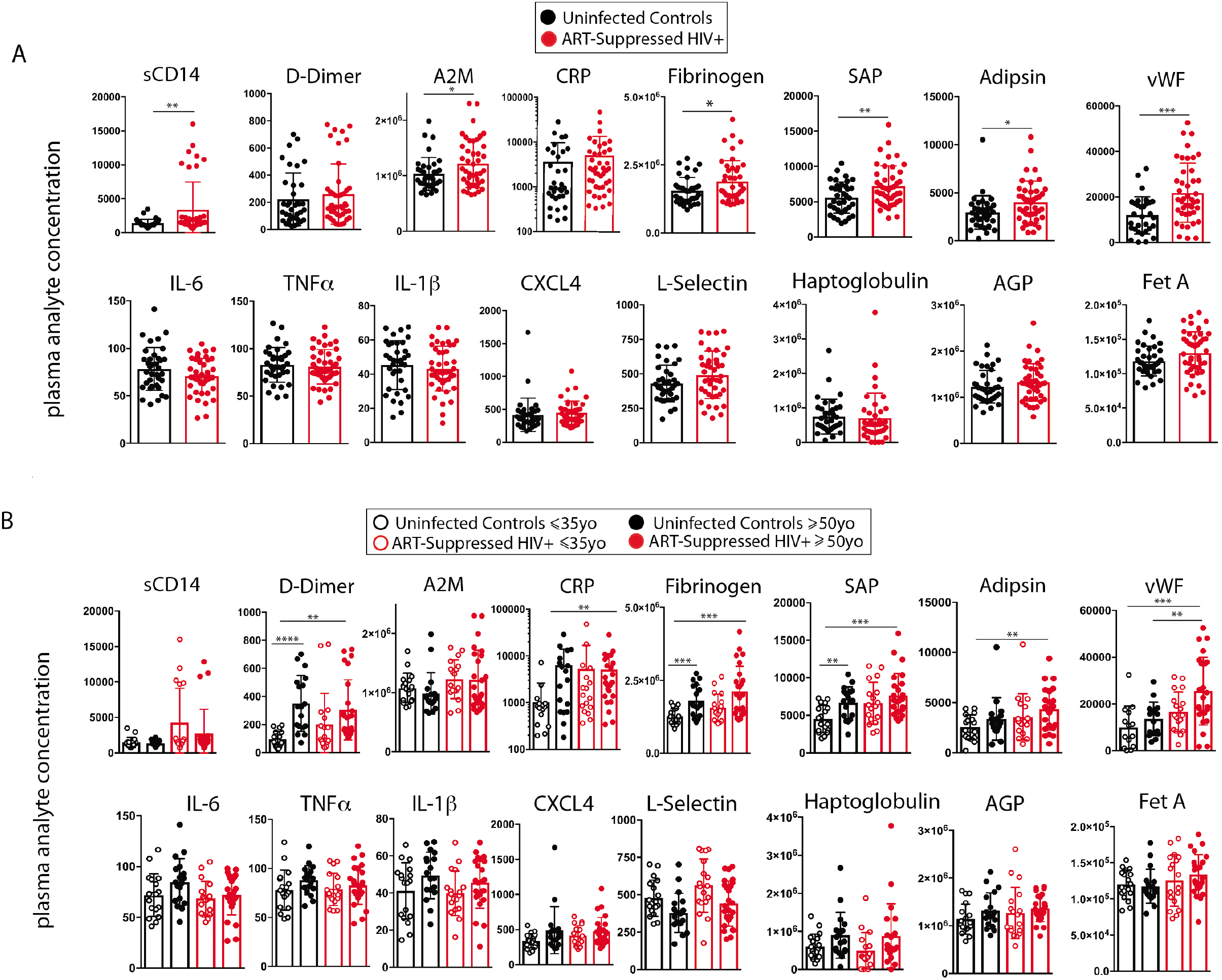
Inflammatory cytokines in plasma in ART-suppressed HIV+ and uninfected controls. 16 plasma analytes were compared between total ART-suppressed HIV+ subjects and uninfected controls (A) and the same subject data with groups further stratified by age (B). Two-tailed T tests were performed for statistical analysis. *p<.05,**p<.01,***p<.001,****p<.0001.unitsare as follows:A2M, CRP, Fetulin A, vWF, Fibrinogen, L-selectin, Haptoglobulin, AGP, Adipsin, CXCL4;ng/ml;Il-6, Il-1β, TNF-αpg/ml;sCD14, D-dimer;ng/ml.

